# Red seaweed (*Asparagopsis taxiformis)* supplementation reduces enteric methane by over 80 percent in beef steers

**DOI:** 10.1101/2020.07.15.204958

**Authors:** B.M. Roque, M. Venegas, R. Kinley, R. deNys, T. L. Neoh, T.L. Duarte, X. Yang, J. K. Salwen, E. Kebreab

**Affiliations:** Department of Animal Science, University of California, Davis, CA 95616, USA; CSIRO Agriculture and Food, Townsville, QLD, AUS; College of Science and Engineering, James Cook University, Townsville, QLD, AUS; Blue Ocean Barns, Redwood City, CA, USA

**Keywords:** *Asparagopsis*, beef cattle, bromoform, enteric methane, greenhouse gas, seaweed

## Abstract

The red macroalgae (seaweed) *Asparagopsis spp.* has shown to reduce ruminant enteric methane (CH_4_) production up to 99% *in vitro.* The objective of this study was to determine the effect of *Asparagopsis taxiformis* on CH_4_ production (g/day per animal), CH_4_ yield (g CH_4_/kg dry matter intake (DMI)), average daily gain (ADG), feed conversion efficiency (FCE), and carcass and meat quality in growing beef steers. Twenty-one Angus-Hereford beef steers were randomly allocated to one of three treatment groups: 0% (Control), 0.25% (Low Dose; LD), and 0.5% (High Dose; HD) *A. taxiformis* inclusion based on organic matter intake. Steers were fed 3 diets: high, medium, and low forage total mixed ration (TMR) representing typical life-stage diets of growing beef steers. The LD and HD treatments over 147 days reduced enteric CH_4_ yield 45 and 68%, respectively; however, there was an interaction between TMR type and the magnitude of CH_4_ yield reduction. Supplementing the low forage TMR reduced CH_4_ yield 69.8% (*P* <0.001) for LD and 80% (*P* <0.001) for HD treatment. Hydrogen (H_2_) yield (g H_2_/DMI) increased significantly (*P*<0.001) 336 and 590% compared to Control for the LD and HD treatments, respectively. No differences were found in carbon dioxide (CO_2_) yield (g CO_2_/DMI), ADG, carcass quality, strip loin proximate analysis and shear force, or consumer taste preferences. DMI tended (*P* = 0.08) to decrease 8% in steers in LD treatment but significantly (*P* = 0.002) reduced 14% in steers in HD treatment. Conversely, FCE tended to increase 7% in steers in LD treatment (*P* = 0.06) and increased 14% in steers in HD (*P* < 0.01) treatment compared to Control. The persistent reduction of CH_4_ by *A. taxiformis* supplementation suggests that this is a viable feed additive to significantly decrease the carbon footprint of ruminant livestock and potentially increase production efficiency.

## 1 INTRODUCTION

Livestock production, particularly ruminants, contributes to anthropogenic greenhouse gas (GHG) emissions globally. These emissions are estimated to be 7.1 Gt carbon dioxide (CO_2_) equivalents annually, approximately 14.5% of the global anthropogenic GHG emissions (Gerber et al., 2013). The majority of GHG emissions from livestock production is mainly in the form of methane (CH_4_), which is produced largely through enteric fermentation and to a lesser extent manure decomposition. Enteric CH_4_ is a natural by-product of microbial fermentation of feed in the digestive tract especially in the rumen. Enteric CH_4_ emissions not only contribute to GHG but also represent an energy loss amounting up to 11% of dietary energy consumption (Moraes et al., 2014). Therefore, reducing enteric CH_4_ emissions contributes to the alleviation of climate change through reduction of GHG emissions from agriculture and can improve productivity through conservation of feed energy otherwise lost as CH_4_. There is potential for mitigation of enteric CH_4_ emissions through a variety of approaches with a focus on the use of feed additives, dietary manipulation and forage quality improvement (Hristov et al., 2013).

Feed additives used in CH_4_ mitigation can either modify the rumen environment or directly inhibit methanogenesis resulting in lower enteric CH_4_ production (g/day per animal) and yield (g/kg dry matter intake [DMI]). Reductions in CH_4_ production of beef cattle through the inhibition of methanogenesis have been reported for feed additives at 22, 93, and 98% for short-chain nitro-compounds (3-nitrooxypropanol; 3-NOP; Dijkstra et al. 2018), synthetic halogenated compounds (Tomkins et al. 2009), and naturally synthesized halogenated compounds in seaweed (Kinley et al. 2020), respectively. The compound 3-NOP inhibits the enzyme methyl-coenzyme M reductase which catalyzes the final step in methanogenesis in rumen archaea (Duin et al. 2016). Halogenated CH_4_ analogs, such as bromoform, act on the same methanogenesis pathway, but do so by binding and sequestering the prosthetic group required by methyl-coenzyme M reductase (Smith et al. 1962; Wood et al, 1968; Johnson et al., 1972). Some haloalkanes are structural analogs of CH_4_, and therefore competitively inhibit the methyl transfer reactions that are necessary in CH_4_ biosynthesis (Ermler et al., 1997; Liu et al., 2011). These CH_4_ analogues include bromochloromethane (BCM), bromoform, and chloroform and have been proven to be the most effective feed additives for reducing CH_4_ production. A 93% reduction of CH_4_ was shown in Brahman cattle with a feed inclusion of BCM at 0.30 g/100 kg LW twice daily for 28 days, however, there was no improvements on feed intake, weight gain, carcass quality or feed efficiency (Tomkins et al. 2009). Conversely, Abecia (2012) reported that the inclusion of BCM at 0.30 g/100 kg once per day decreased CH_4_ production 33% and increased milk production 36%. The authors speculated that increased milk production in BCM treated cows could be attributed to a shift to more propionate production in the rumen, which is a hydrogen (H_2_) sink and provides more energy compared to other volatile fatty acids. However, long-term efficacy of CH_4_ analogues in the rumen remains to be confirmed. Tomkins et al. (2009), for example, reported a second experiment resulting in a 57.6% CH_4_ reduction after 30 days of treatment which is far less than the reductions found during the first 28 days. Additionally, chloroform fed to fistulated dairy cows was effective at reducing enteric CH_4_ production through reduced abundance and activity of methanogenic archaea, but only over a 42 day period (Knight, 2011).

Types of feedstuffs can also drive CH_4_ production by providing different substrates to microbial populations which are the drivers of volatile fatty acid (VFA) production in the rumen. There are ways to influence the types of VFA produced in the rumen by changing the types of feed in the diet (Russell and Wallace, 1997, Van Soest, 1994). This is important for two reasons; first VFA represent the amount of energy available to the animal for means of animal productivity and second VFA pathways, such as the production of propionate, are able to utilize reducing equivalents that normally would be shifted to methanogenesis (Blaxter and Clappteron, 1965, Johnson and Johnson, 1995). Concentrates contain non-structural carbohydrates, such as starch and sugar, that are rapidly fermented which drives pH down, which negatively impact methanogenic populations, and are an effective way to increase propionate production (Bannink et al., 2006, 2008). Forages contain structural carbohydrates, such as neutral detergent fiber (NDF), and have been linked to CH_4_ production (Niu et al, 2018). As NDF in diet increases, rumen pH also increases resulting in preferential production of acetate over propionate, which generates reducing equivalents such as H_2_ that is shifted toward methanogenesis (Hungate, 1966, Janssen, 2010). Not only can NDF play a significant role in CH_4_ production, it has also been suggested to impact the efficacy of anti-methanogenic compounds added to feed (Dijkstra et al. 2018).

Red seaweeds, particularly the genus *Asparagopsis*, are considered potent anti-methanogenic organisms due to their capacity to synthesize and encapsulate halogenated CH_4_ analogues such as bromoform and dibromochloromethane within specialized gland cells as a natural defense mechanism against predation (Paul et al., 2006). Machado et al. (2014) compared a diversity of tropical macroalgae, including freshwater and marine species, and found that *Asparagopsis taxiformis* at 17% of OM had the largest reduction of CH_4_ production *in vitro* with a 98.9% average reduction. In the stepwise progression of evaluating the seaweeds Kinley et al. (2016a; 2016b) explored reduced inclusion rates to determine the *in vitro* minimum effective inclusion level of 2% of OM. In this process only *A. taxiformis* retained anti-methanogenic capability at a very low inclusion. *A. taxiformis* reduced CH_4_ production more effectively than synthetic halogenated CH_4_ analogs at equivalent concentrations *in vitro*, largely due to multiple anti-methanogenic bioactives working synergistically (Machado et al., 2018). Importantly, the concentration of bioactives in *Asparagopsis spp*, in particular bromoform, has a significant effect on the reduction of CH_4_ in animal trials. For example, Roque et al (2019b) tested the effects of the seaweed *A. armata* (1.3 mg bromoform/g DM) *in vivo* fed to dairy cattle for a two week duration and reported up to 67% reduction of enteric CH_4_ production using a 1% seaweed inclusion rate on organic matter (OM) basis in a total mixed ration (TMR). Kinley et al. (2020) reported up to 98% CH_4_ reduction using 0.2% of OM inclusion of *A. taxiformis* (6.6 mg bromoform/g DM) during a 90-day feeding regime typical of feedlot TMR. These studies confirm that seaweed quality measured as concentration of bioactive at feeding and the basal diet formulation have an impact on the efficacy of the seaweed, and that there is heightened response *in vivo* compared to *in vitro*.

For adoption of the seaweed by industry it is crucial that meat quality be maintained or improved. As with any feed additive, feeding *A. taxiformis* to livestock has the potential for changes in meat quality, tenderness and taste, and consumer acceptability. Marbling, for instance, directly impacts flavor and juiciness and it has been shown that marbling can directly influence consumer preference with some willing to pay a premium (Killinger et. al., 2004).

Therefore, the objectives of this study were to (1) measure the long-term effects of *A. taxiformis* over 147 day period, (2) test the efficacy of CH_4_ reductions of supplementing *A. taxiformis* to high, medium, and low forage basal TMRs, (3) quantify the effects of *A. taxiformis* supplementation on production parameters, meat quality (including taste), and bromoform residues within the meat and liver.

## 2 MATERIALS AND METHODS

This study was approved by the Institutional Animal Care and Use Committee of the University of California, Davis (Protocol No. 20803).

### 2.1 Study design, animals, and diets

Twenty-one Angus-Hereford cross beef steers, blocked by weight, were randomly allocated to one of three treatment groups: 0% (Control), 0.25% (Low Dose; LD), and 0.5% (High Dose; HD) inclusion rates of *A. taxiformis* based on OM intake. One steer on HD treatment was injured during the last 3 weeks of the trial and all data from this steer was removed from statistical analysis. The experiment followed a completely randomized design, with a 2-week covariate period before treatment began followed by 3-week data collection intervals for 21-weeks; a total of 147 days of seaweed treatment (Figure 1). During data collection intervals, alfalfa pellets offered through the CH_4_ measuring device (GreenFeed system, C-Lock, Inc., Rapid City, SD) were included as part of daily feed intake. The steers were individually housed and were approximately 8 months of age with an average BW of 352 ± 9 kg at induction to the trial. Steers were fed 3 diets during the study; high (starter diet), medium (transition diet), and low (finisher diet) forage TMRs, which are typical life-stage TMRs of growing beef steers (Table 1). Samples from the three diets and alfalfa pellets were collected once a week and bags of *A. taxiformis* were randomly sampled and analyzed (Table 2) for dry matter, acid detergent fiber, NDF, lignin, starch, crude fat, total digestible nutrient and mineral content (Cumberland Valley Analytical Services, Waynesboro, PA). Steers were offered water *ad libitum*.

**TABLE 1.** Ingredients of the experimental diet containing high, medium, and low forage concentrations (% of DM)

| Ingredients | High forage | Medium forage | Low forage |
| --- | --- | --- | --- |
| *Forage* |  |  |  |
| Alfalfa hay | 35.0 | 25.0 | 5.00 |
| Wheat hay | 25.0 | 15.0 | 6.00 |
| Dry distillers grain | 12.0 | 14.0 | 6.00 |
| *Concentrate* |  |  |  |
| Rolled corn | 20.0 | 37.0 | 72.0 |
| Molasses | 5.0 | 5.00 | 3.00 |
| Fat | 1.5 | 2.00 | 3.00 |
| Urea | 0.35 | 0.40 | 1.80 |
| Beef trace salt | 0.32 | 0.32 | 1.00 |
| Calcium carbonate | 0.82 | 1.15 | 1.80 |
| Magnesium oxide |  |  | 0.20 |
| Potassium chloride |  |  | 0.50 |

**TABLE 2.** Nutritional composition of experimental diets, *Asparagopsis taxiformis*, and alfalfa pellets

|  | High forage | Medium forage | Low forage | Pellets | *A. taxiformis* |
| --- | --- | --- | --- | --- | --- |
| *% Dry matter* |  |  |  |  |  |
| Organic matter | 92.1 | 93.1 | 94.8 | 88.6 | 50.9 |
| Crude protein | 17.2 | 17.4 | 13.2 | 17.1 | 16.8 |
| ADF | 22.6 | 16.7 | 10.5 | 28.1 | 11.5 |
| NDF | 33.1 | 25.8 | 18.6 | 35.9 | 33.7 |
| Lignin | 4.08 | 3.05 | 1.73 | 6.16 | 4.08 |
| Starch | 16.9 | 25.0 | 46.7 | 0.90 | 0.35 |
| Crude fat | 4.92 | 6.04 | 6.77 | 3.02 | 0.63 |
| Calcium | 0.77 | 1.00 | 0.50 | 2.06 | 5.29 |
| Phosphorus | 0.33 | 0.38 | 0.28 | 0.24 | 0.18 |
| Magnesium | 0.38 | 0.38 | 0.23 | 0.37 | 0.81 |
| Potassium | 1.74 | 1.42 | 0.94 | 2.10 | 2.02 |
| Sodium | 0.18 | 0.25 | 0.30 | 0.20 | 6.34 |
| *Parts per million* |  |  |  |  |  |
| Iron | 438 | 335 | 127 | 1508 | 8494 |
| Manganese | 61.7 | 56.0 | 64.0 | 88.0 | 142.5 |
| Zinc | 43.2 | 51.50 | 58.0 | 19.0 | 53.5 |
| Copper | 8.67 | 8.00 | 7.00 | 10.0 | 22.5 |

**FIGURE 1.**
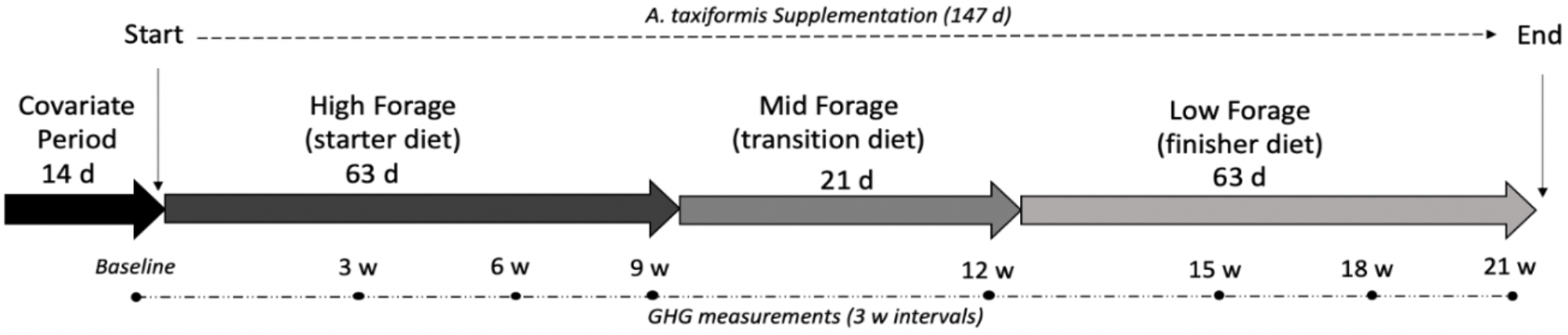
Experimental timeline including covariate period, total days for *Asparagopsis taxiformis* supplementation, dietary regime, days on diets, and greenhouse gas measurement intervals.

The *A. taxiformis* used as a feed additive was provided by Commonwealth Scientific and Industrial Research Organization (CSIRO) Australia. The seaweed was collected during the gametophyte phase from Humpy Island, Keppel Bay, QLD (23°13’01”S, 150°54’01”E) by Center for Macroalgal Resources and Biotechnology of James Cook University, Townsville, Queensland, Australia. Once collected, it was frozen and stored at −15 °C then freeze dried at Forager Food Co., Red Hills, Tasmania, Australia and later ground to a size of 2–3 mm. Total seaweed inclusion ranged from 46.7 to 55.7 g/day for LD and 76.1 to 99.4 g/day for HD treatment. The seaweed used in the study contained bromoform at a concentration of 7.8 mg/g dry weight as determined by Bigelow Analytical Services, East Boothbay, ME, USA. To increase palatability and adhesion to feed, 200 ml of molasses and 200 ml of water was added to *A. taxiformis* then hand mixed into the total mixed ration for each animal. The Control group also received 200 ml of both molasses and water with their daily feed to ensure *A. taxiformis* was the only difference between the three treatments. Steers were fed 105% of the previous day’s intake twice daily at 0600 and 1800 hours. Daily feed intake was calculated as the TMR offered minus feed refusal weights.

### 2.2 Sample collection and analysis

Methane, CO_2_, and H_2_ gas emissions from steers were measured using the GreenFeed system (C-Lock Inc., Rapid City, SD, USA). Gas emissions were measured during the covariate period and experimental period during weeks 3, 6, 9, 12, 15, 18, and 21. In each measurement period, gas emission data were collected during 3 consecutive days as follows: starting at 07:00, 13:00, and 1900 hours (sampling day 1); 0100, 1000, and 1600 hours (sampling day 2); and 2200 and 0400 hours (sampling day 3). Breath gas samples were collected for 3 to 5 minutes followed by a 2-minute background gas sample collection. The GreenFeed system was calibrated before each measurement period with a standard gas mixture containing (mol %): 5000 ppm CO_2_, 500 ppm CH_4_, 10 ppm H_2_, 21% O_2_ and nitrogen as a balance (Air Liquide America Specialty Gases, Rancho Cucamonga, CA). Recovery rates for CO_2_ and CH_4_ observed in this study were within +/− 3% of the known quantities of gas released. Alfalfa pellets were offered at each sampling event as bait feed and was kept below 10% of the total DMI during each 3 day measurement period. The composition of alfalfa pellets is shown in Table 2. Feed intake and feed costs were recorded daily and bodyweight (BW) was measured weekly.

After the feeding trial was completed, all 20 steers were sent to a USDA-inspected commercial packing plant for slaughter. On the day of slaughter, steers were marked and followed throughout the process. On the first day, livers were collected and stored on dry ice until placed in a −20°C freezer. Carcasses were aged for 48 hours in a large cooler and then graded by a certified USDA grader. Directly after grading, carcasses were sent to fabrication where the strip loin from the left side of each carcass was cut and saved for further analysis. All 20 strip loins were placed on ice and transported back to the University of California, Davis where they were cryovac packaged and stored at 4°C in dark for 14 days. After 14-day of aging, strip loins were cut into steaks (2.45 cm thickness) and individually vacuum packaged and stored at −20°C. Steaks and livers were analyzed for bromoform concentrations using Shimadzu QP2010 Ultra GC/MS following a modified protocol described by Paul et al. (2006) (Bigelow Analytical Services, East Boothbay, ME, USA). The limits of bromoform detection and quantification were 0.06 mg/kg and 0.20 mg/kg, respectively.

To test for objective tenderness, slice shear force (SSF) and Warner-Brazler shear force (WBSF) were measured following the protocol described by AMSA (2016). One steak from each animal was thawed overnight and cooked to an internal temperature of 71°C. Within 1 to 2 minutes after cooking, the SSF were measured using machine texture analyzer (TMS Pro Texture Analyzer, Food Technology corporation, Sterling, VA, USA) with a crosshead at the speed of 500 mm/minute. To test WBSF, cooked steaks were cooled at 4°C overnight, and then four cores were cut using WEN 8-inch 5 Speed Drill Press from one steak from each animal parallel to the muscle fiber orientation. The WBSF was measured using the TMS Pro texture analyzer with a Warner Bratzler blade (2.8 mm wide) and a crosshead at speed of 250 mm/minute. The average peak forces for all four cores were recorded.

A consumer sensory panel was conducted at UC-Davis. Strip steaks were thawed at 4°C for 24 hours then cooked to an internal temperature of 71°C using a George Foreman clamshell (Spectrum Brands, Middleton, WS, USA). Internal temperature was taken from the geometric center of each steak using a K thermocouple thermometer (AccuTuff 351, model 35100, Cooper-Atkins Corporation, Middlefield, CT, USA). Following cooking, steaks were rested for 3 minutes then cut into 1.5 cm^3^ pieces. Each steak was randomly assigned a unique three digit number, placed into glass bowls covered in tin foil then stored in an insulated food warmer (Carlisle model PC300N03, Oklahoma, OK, USA) for longer than 30 minutes prior to the start of each sensory evaluation session. A total of 112 participants evaluated steak samples during one of the 5 sessions held over a 4-day period. Each participant evaluated a total of three steak samples, one from each treatment group, with a minimum of two 1.5 cm^3^ pieces per steak. Each participant was asked to evaluate tenderness, flavor, juiciness, and overall acceptance using a 9-point hedonic scale (1 = Dislike extremely and 9 = Like extremely).

### 2.3 Statistical analysis

Statistical analysis was performed using R statistical software (version 3.6.1; The R Foundation for Statistical Computing, Vienna, Austria). The linear mixed-effects models (lme) procedure was used with the steer as the experimental unit. GreenFeed emission data were averaged per steer and gas measurement period, which was then used in the statistical analysis. Dry matter intake and cost per kg of gain (CPG) data was averaged by week and used in the statistical analysis. Average daily gain (ADG) was calculated by subtracting initial BW from final BW then dividing by the number of experimental days for each diet regimen and the duration of the study (i.e. 63 days on high forage (starter) TMR, 21 days on medium forage (transition) TMR, then 63 days on low forage (finisher) TMR with total study duration of 147 days). Feed conversion efficiency (FCE) was calculated by dividing ADG by DMI for each diet regimen and the duration of the study.

The statistical model included treatment, diet, treatment × diet interactions, and the covariate term, with the error term assumed to be normally distributed with mean = 0 and constant variance. Individual animal was used as random effect, whereas all other factors were considered fixed. Data was analyzed as repeated measures with an autoregressive 1 correlation structure. Statistical significance was established when *P* ≤ 0.05 and a trend at 0.05 > *P* ≤ 0.10. The consumer sensory evaluation data were analyzed using the Kruskal-Wallis test. The Dunn’s test with *P*-value adjustment following Bonferroni methods was used for post-hoc pair-wise comparisons.

## 3 RESULTS

### 3.1 Gas parameters

The emissions as production (g/day per animal) and yield (g/DMI kg) of CH_4_, H_2_, and CO_2_ gases from the steers in the three treatment groups (Control, LD, HD) are presented in Figure 2 (for the duration of the trial) and Table 3 (divided by the three diet regimes). Where *P* values for significant effects are not given, they are *P* < 0.01. Inclusion of *A. taxiformis* in the TMR had a significant linear reduction in enteric CH_4_ production and yield. Methane production for the collective feeding stages of experimental period resulted in a reduction of 50.6 and 74.9% for LD and HD treatments, respectively, compared to Control. Methane yields for LD and HD were 45 and 68% lower, respectively, compared to Control. However, there was an interaction between diet formulation and magnitude of CH_4_ production and yield reduction. Methane production in steers on the high forage TMR with *A. taxiformis* inclusion was significantly reduced 36.4% for LD and 58.7% for HD and CH_4_ yield was significantly reduced 32.7% for LD and 51.9% for HD compared to the Control steers. Methane production during the medium forage TMR phase was significantly reduced 51.8 and 86.8%, for LD and HD treatments compared to Control, respectively, whereas CH_4_ yield was significantly reduced 44.6 and 79.7%, respectively. Steers fed low forage TMR in LD and HD treatments produced 72.4 and 81.9% lower CH_4_ compared to Control, respectively. Similarly, their CH_4_ yield was significantly reduced 69.8 for LD and 80.0% for HD.

**TABLE 3.**
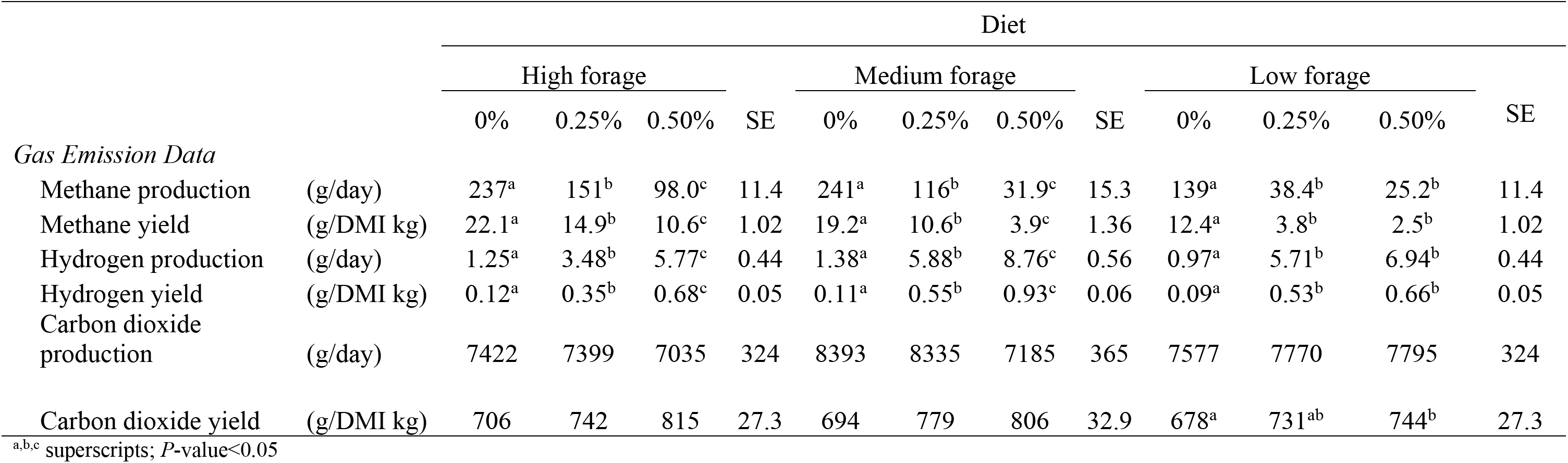
Effect of *Asparagopsis taxiformis* inclusion levels of 0%, 0.25%, and 0.5% of feed organic matter to three stages of beef cattle diets on gas parameters

|  |  | Diet |  |  |  |  |  |  |  |  |  |  |  |
| --- | --- | --- | --- | --- | --- | --- | --- | --- | --- | --- | --- | --- | --- |
|  |  | High forage |  |  |  | Medium forage |  |  |  | Low forage |  |  |  |
|  |  | 0% | 0.25% | 0.50% | SE | 0% | 0.25% | 0.50% | SE | 0% | 0.25% | 0.50% | SE |
| *Gas Emission Data* |  |  |  |  |  |  |  |  |  |  |  |  |  |
| Methane production | (g/day) | 237<sup>a</sup> | 151<sup>b</sup> | 98.0<sup>c</sup> | 11.4 | 241<sup>a</sup> | 116<sup>b</sup> | 31.9<sup>c</sup> | 15.3 | 139<sup>a</sup> | 38.4<sup>b</sup> | 25.2<sup>b</sup> | 11.4 |
| Methane yield | (g/DMI kg) | 22.1<sup>a</sup> | 14.9<sup>b</sup> | 10.6<sup>c</sup> | 1.02 | 19.2<sup>a</sup> | 10.6<sup>b</sup> | 3.9<sup>c</sup> | 1.36 | 12.4<sup>a</sup> | 3.8<sup>b</sup> | 2.5<sup>b</sup> | 1.02 |
| Hydrogen production | (g/day) | 1.25<sup>a</sup> | 3.48<sup>b</sup> | 5.77<sup>c</sup> | 0.44 | 1.38<sup>a</sup> | 5.88<sup>b</sup> | 8.76<sup>c</sup> | 0.56 | 0.97<sup>a</sup> | 5.71<sup>b</sup> | 6.94<sup>b</sup> | 0.44 |
| Hydrogen yield | (g/DMI kg) | 0.12<sup>a</sup> | 0.35<sup>b</sup> | 0.68<sup>c</sup> | 0.05 | 0.11<sup>a</sup> | 0.55<sup>b</sup> | 0.93<sup>c</sup> | 0.06 | 0.09<sup>a</sup> | 0.53<sup>b</sup> | 0.66<sup>b</sup> | 0.05 |
| Carbon dioxide production | (g/day) | 7422 | 7399 | 7035 | 324 | 8393 | 8335 | 7185 | 365 | 7577 | 7770 | 7795 | 324 |
| Carbon dioxide yield | (g/DMI kg) | 706 | 742 | 815 | 27.3 | 694 | 779 | 806 | 32.9 | 678<sup>a</sup> | 731<sup>ab</sup> | 744<sup>b</sup> | 27.3 |

<sup>a,b,c</sup> superscripts; *P*-value<0.05

**FIGURE 2.**
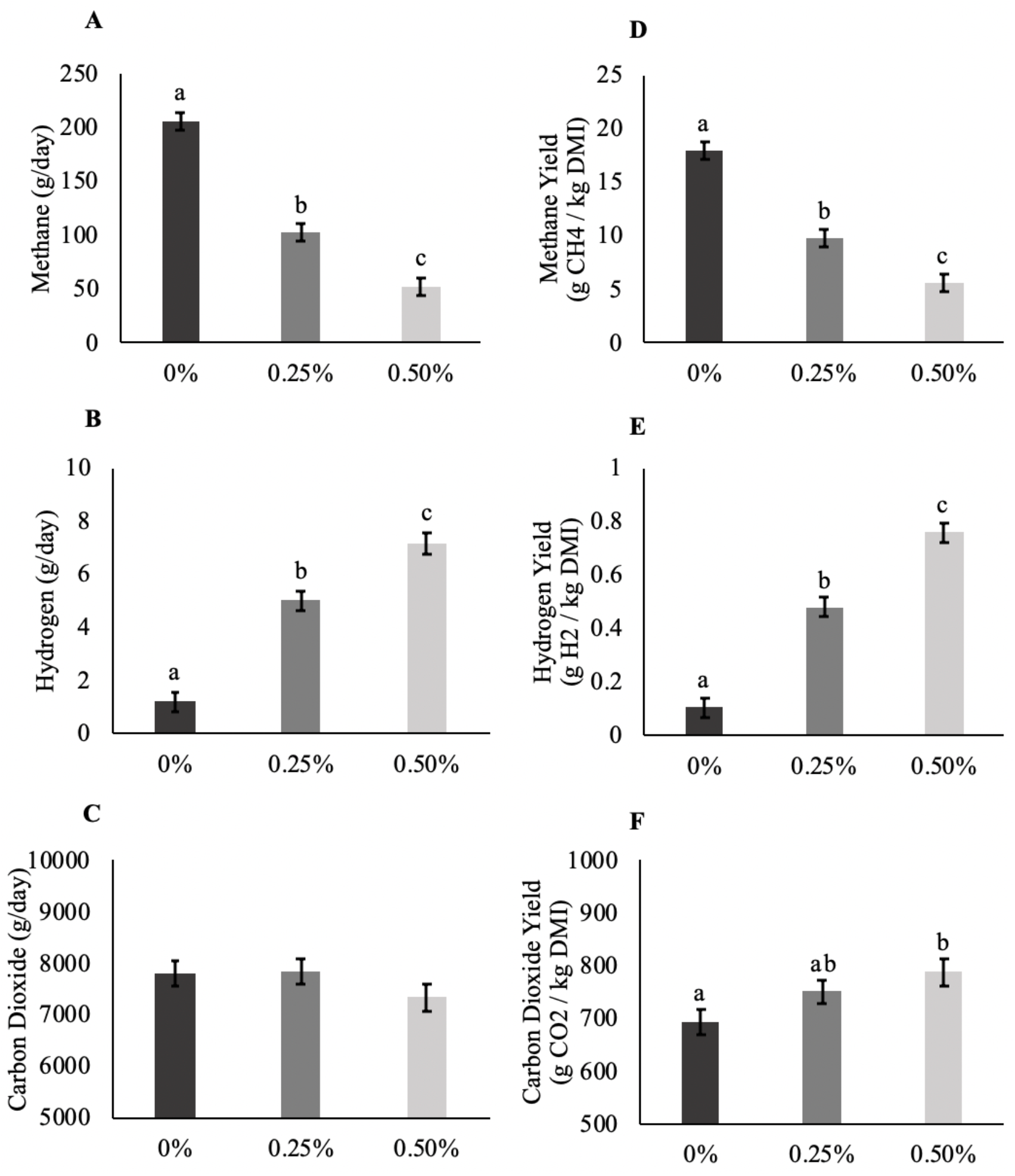
Means, standard deviations, and statistical differences of methane, hydrogen, and carbon dioxide production (g/d) (A,B,C), and yield (g/kg dry matter intake (DMI)) (D,E,F) for 0%, 0.25%, and 0.50% *Asparagopsis taxiformis* inclusion. Means within a graph with different alphabets differ (*P* < 0.05)

Hydrogen production significantly increased 318 and 497% and H_2_ yield also increased significantly 336 and 590% compared to Control for the LD and HD treatments for the collective feeding stages of the experiment, respectively. Hydrogen production in steers receiving *A. taxiformis* to high forage TMR in the LD and HD treatments significantly increased 177 (*P* = 0.03) and 360%, respectively. Hydrogen yield increased 198 (*P* = 0.04) and 478% in steers on LD and HD treatments, respectively. Hydrogen production from steers on the medium forage TMR increased 326 (*P* = 0.03) and 535% (*P* < 0.01), for LD and HD treatments, respectively, whereas H_2_ yield significantly increased 404 and 753%, respectively. Supplementation of *A. taxiformis* to the low forage TMR fed to steers significantly increased H_2_ production 419 and 618% and reduced H_2_ yield 503 and 649% in LD and HD treatments, respectively. Carbon dioxide (CO_2_) production was not affected by either LD or HD treatments compared to Control. However, CO_2_ yield was significantly greater in HD group compared to Control (*P* = 0.03).

### 3.2 Animal production parameters

Dry matter intake, ADG, feed conversion efficiency (ADG/DMI; FCE) and cost per gain ($USD/kg weight gain; CPG) as impacted by treatment groups (Control, LD, HD) for the collective feeding stages is presented in Table 4 and for the individual stages and TMRs in Table 5. Initial BW, final BW, carcass weight and total weight gained are shown in Table 4. Considering all feeding stages, steers on the LD treatment tended (*P* = 0.08) to decrease their DMI 8% and was significantly reduced 14% in steers on the HD (*P* < 0.01) treatment compared to Control. Steers fed the high and medium forage TMR in the HD treatment decreased their DMI 18.5 (*P* = 0.01) and 18.0% (*P* < 0.01), respectively, compared to Control. No significant effects were observed in ADG by the LD or HD treatment groups. With the reduction of DMI in LD and HD treatments and similar ADG among all 3 treatments, FCE tended to increase 7% in LD (*P* = 0.06) treatment and significantly increased 14% in steers in HD (*P* < 0.01) treatment. When averaged throughout the experiment as well as within the 3 TMR stages, CPG was not statistically significant. However, over the collective feeding stages, CPG was consistently lower in HD and LD groups compared to Control with approximately $0.37 USD/kg gain differential between HD and Control and $0.18 USD/kg gain differential between LD and Control. Additionally, cost differentials for HD were $0.29, $0.40, and $0.34 USD/kg gain and for LD were $0.15, $0.49, and $0.34 USD/kg gain for the high, medium, and low forage TMRs, respectively.

**TABLE 4.** Effect of *Asparagopsis taxiformis* inclusion levels of 0%, 0.25%, and 0.5% of feed organic matter on animal parameters over 21 weeks

|  |  | 0% | 0.25% | 0.50% | SE |
| --- | --- | --- | --- | --- | --- |
| *Animal Parameters* |  |  |  |  |  |
| Dry matter intake | (kg/day) | 11.32<sup>a</sup> | 10.42<sup>ab</sup> | 9.69<sup>b</sup> | 0.29 |
| Average daily gain | (kg/day) | 1.60 | 1.52 | 1.56 | 0.06 |
| Feed conversion efficiency | (ADG/DMI) | 0.14<sup>a</sup> | 0.15<sup>a</sup> | 0.16<sup>b</sup> | 0.01 |
| Cost per gain | (\$/kg gain) | 2.25 | 2.07 | 1.88 | 0.18 |
| Initial body weight | (kg) | 357 | 348 | 350 | 9.21 |
| Final body weight | (kg) | 589 | 572 | 587 | 11.1 |
| Carcass weight | (kg) | 370 | 361 | 350 | 13.4 |
| Total gain | (kg) | 232 | 224 | 236 | 6.09 |

<sup>a,b,c</sup> superscripts; $P$ -value <0.05

**TABLE 5.** Effect of *Asparagopsis taxiformis* inclusion levels of 0%, 0.25%, and 0.5% feed organic matter to beef cattle diets on animal parameters

|  |  | Diet |  |  |  |  |  |  |  |  |  |  |  |
| --- | --- | --- | --- | --- | --- | --- | --- | --- | --- | --- | --- | --- | --- |
|  |  | High forage |  |  |  | Medium forage |  |  |  | Low forage |  |  |  |
|  |  | 0% | 0.25% | 0.50% | SE | 0% | 0.25% | 0.50% | SE | 0% | 0.25% | 0.50% | SE |
| *Animal Parameters* * |  |  |  |  |  |  |  |  |  |  |  |  |  |
| Dry matter intake | (kg/day) | 10.31<sup>a</sup> | 9.69<sup>ab</sup> | 8.40<sup>b</sup> | 0.33 | 12.18<sup>a</sup> | 10.81<sup>ab</sup> | 9.99<sup>b</sup> | 0.33 | 11.47<sup>a</sup> | 10.77<sup>ab</sup> | 10.67<sup>b</sup> | 0.33 |
| Average daily gain | (kg/day) | 1.58 | 1.61 | 1.53 | 0.11 | 1.62 | 1.50 | 1.38 | 0.11 | 1.60 | 1.44 | 1.75 | 0.11 |
| Feed conversion | (ADG/DMI) | 0.15 | 0.17 | 0.18 | 0.01 | 0.13 | 0.14 | 0.14 | 0.01 | 0.14 | 0.13 | 0.17 | 0.01 |
| Cost per gain | ( \$/kg ) | 1.83 | 1.68 | 1.54 | 0.25 | 2.67 | 2.18 | 2.20 | 0.26 | 2.24 | 2.35 | 1.90 | 0.27 |

<sup>a,b,c</sup> superscripts; *P*-value<0.05

### 3.3 Carcass quality parameters

There was no statistical difference between treatment groups for rib eye area (Table 6). No effects were found between Control, LD, and HD in moisture, protein, fat, ash, carbohydrates, or calorie content of strip loins (Table 6). The average WBSF values for the Control, LD and HD groups were 2.81, 2.66 and 2.61 kg, respectively. Additionally, the SSF averages were measured as 17.1 for Control, 16.75 for LD and 17.4 kg for HD treatments. No significant differences (*P* > 0.05) were found in shear force resistance among treatment groups. Mean scores of all sensory attributes (tenderness, juiciness, and flavor) by consumer panels were not significantly different (*P* > 0.05) among treatment groups (Table 6). The taste panel considered all steaks, regardless of treatment group, to be moderately tender as well as moderately juicy. This was consistent with the taste panel stating that they moderately liked the flavor of all steaks regardless of treatment group. There was no difference (*P* > 0.05) in overall acceptability among treatment groups.

**TABLE 6.** Effects of *Asparagopsis taxiformis* supplementation on carcass quality, proximate analysis, shear force, and consumer panel preference.

|  | Level of *Asparagopsis taxiformis* inclusion |  |  |  |
| --- | --- | --- | --- | --- |
|  | 0% | 0.25% | 0.50% | SE |
| *Carcass Quality* |  |  |  |  |
| Rib eye area (inches) | 11.3 | 11.2 | 10.6 | 0.37 |
| *Proximate Analysis* |  |  |  |  |
| Moisture (g/100g) | 53.9 | 55.4 | 55.3 | 1.7 |
| Protein (g/100g) | 16.1 | 17.1 | 17.2 | 0.7 |
| Fat (g/100) | 29.1 | 26.1 | 26.3 | 2.2 |
| Ash (g/100g) | 0.73 | 0.86 | 0.88 | 0.05 |
| Carbohydrates (g/100g) | 0.24 | 1.01 | 0.42 | 0.38 |
| Calories | 327 | 307 | 307 | 17.7 |
| Iodine (PPM) | 0.00<sup>a</sup> | 0.08<sup>b</sup> | 0.15<sup>c</sup> | 0.02 |
| *Shear Force* |  |  |  |  |
| Warner- Bratzler (kgf) | 2.81 | 2.66 | 2.61 | 0.24 |
| Slice Shear Force (kgf) | 17.1 | 16.8 | 17.4 | 1.87 |
| *Consumer Panel<sup>1</sup>* |  |  |  |  |
| Tenderness | 6.72 | 6.68 | 6.45 | 0.17 |
| Juiciness | 6.35 | 6.33 | 6.07 | 0.17 |
| Flavor | 6.63 | 6.34 | 6.24 | 0.15 |
| Overall | 6.66 | 6.36 | 6.46 | 0.16 |

<sup>a,b,c</sup> superscripts; *P*-value <0.05
<sup>1</sup>A 9-point hedonic scale was used
(1 = Dislike extremely and 9 = Like extremely).

There was a linear increase in iodine concentrations in both LD (*P* < 0.01) and HD (*P* < 0.01) compared to Control. Iodine concentrations for the Control treatment group were below detection levels, which was set at 0.10 mg/g (Table 6). However, 5 out of 7 steers in LD treatment group had iodine levels above the detection level with a treatment average of 0.08 mg/g (*P* < 0.01). All 6 steers in the HD treatment group were found to contain iodine levels above the detection level with concentration levels ranging between 0.14 – 0.17 mg/g with a mean of 0.15 mg/g (*P* < 0.01). Bromoform concentrations for all treatment groups were below detection levels, which were 0.06 mg/kg.

## 4 DISCUSSION

### 4.1 Enteric methane production and yield

This study demonstrated that dietary inclusion of *A. taxiformis* induces a consistent and considerable reduction in enteric CH_4_ production from steers on a typical feedlot style diet. Enteric CH_4_ is the largest contributor to GHG emissions from livestock production systems. Similar reductions in CH_4_ yield, which is standardized by DMI, has also been established. There is a concern that feed additives and other CH_4_ reducing agents decrease in efficacy over time (Knight et al., 2011, Rumpler et al., 1986). This study provided evidence that the seaweed inclusion was effective in reducing CH_4_ emissions, which persisted for the duration of the study of 147 days (Figure 3). Notably, until this study the longest exposure to *A. taxiformis* had been demonstrated for steers in a study ending after a 90-d finishing period (Kinley et al. 2020). To date, only three *in vivo* studies have been published using *Asparagopsis spp* to reduce enteric CH_4_ emissions in sheep (Li et al. 2018), lactating dairy cattle (Roque et al., 2019b), and feedlot Brangus steers (Kinley et al., 2020). All studies show considerable if not variable reduction in enteric CH_4_ emissions. The differences in efficacy are likely due to levels of seaweed inclusion, formulation of the diets, and differences in seaweed quality based on bromoform concentrations.

**FIGURE 3.**
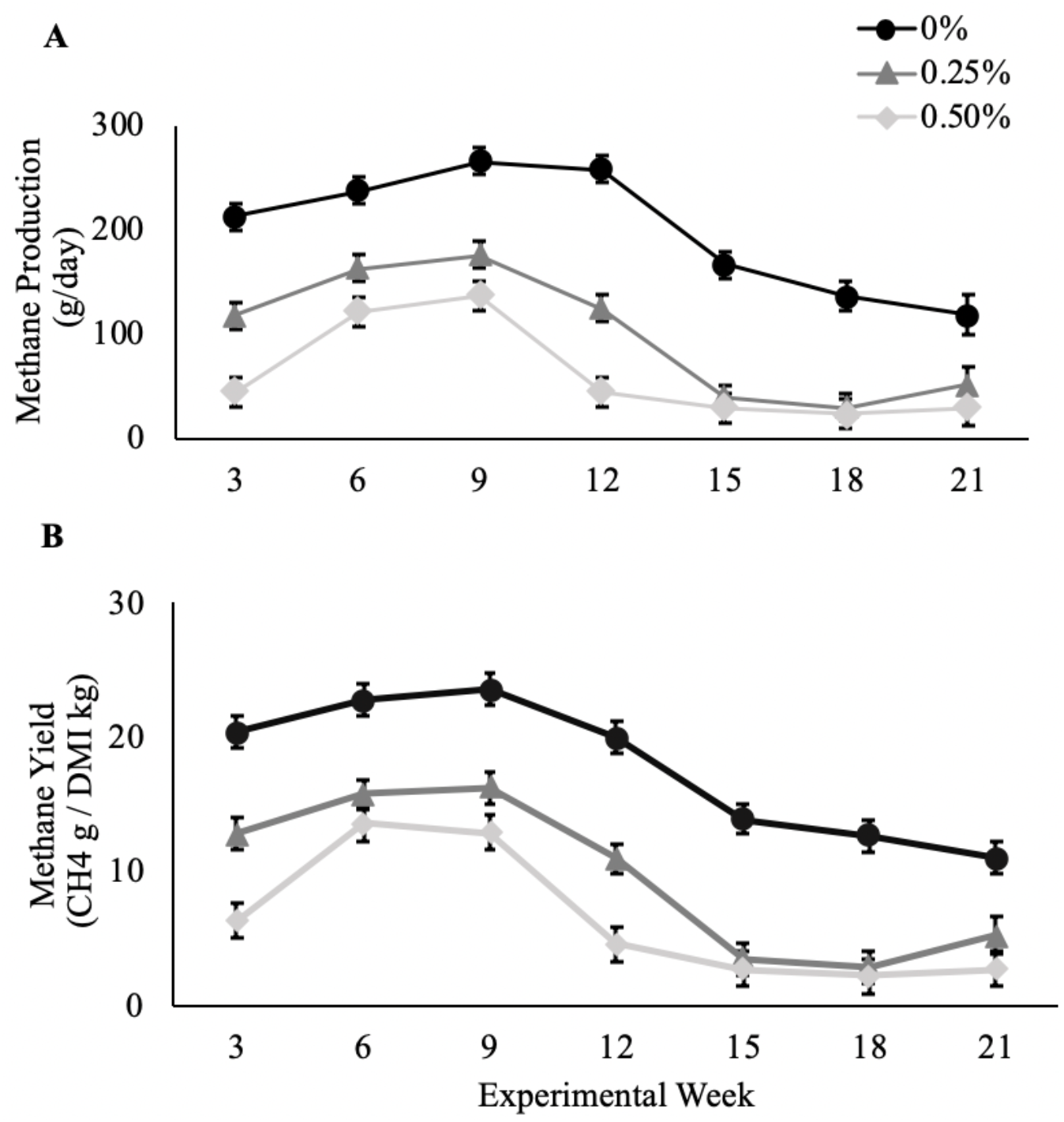
Methane production [g CH_4_/day] (A) and methane yield [g CH_4_/kg DMI] (B) from beef steers supplemented with *Asparagopsis taxiformis* at 0%, 0.25%, and 0.5% of basal total mixed ration on an organic matter basis during the 21 week experimental period. Data points are treatment means for each gas collection timepoint and error bars represent standard errors.

It has been previously hypothesized that NDF levels can also influence the rate at which CH_4_ is reduced with the inclusion of inhibitors (Dijkstra et al. 2018). In the current study, the magnitude of reductions in CH_4_ production were negatively correlated (r^2^ = 0.89) with NDF levels in the 3 diet regimens that contained 33.1% (high forage), 25.8% (medium forage), and 18.6% (low forage) NDF levels. Enteric CH_4_ production was reduced 32.7, 44.6 and 69.8% in steers on the LD treatment and 51.9, 79.7, and 80.0% on HD treatment with high, medium and low forage TMRs, respectively. The low forage TMR, containing the lowest NDF levels, was the most sensitive to the inclusion of *A. taxiformis* with CH_4_ reductions above 70% at equivalent inclusion levels compared to the higher forage TMRs. Li et al. (2018) reported an 80.6% reduction of CH_4_ yield in sheep fed diets containing 55.6% NDF, however, the level of *A. taxiformis* intake by the sheep was unclear but was offered at 6 times greater levels than the HD treatment in our study. Roque et al. (2019b) showed 42.7% reduction in CH_4_ yield in lactating dairy cattle fed a diet containing 30.1% NDF at 1% inclusion rate of *A. armata*. The high forage TMR in our study had a similar NDF level to Roque et al. (2019b), however, had approximately double the reduction of CH_4_, even when consuming 50% less seaweed. These differences relate to a large degree to the quality of seaweed in terms of the concentration of bromoform, which was 1.32 mg/g in Roque et al. (2019b) compared to 7.82 mg/g in the current study. Kinley et al. (2020) conducted an *in vivo* study focused on feedlot steers using the same collection of *A. taxiformis* as sub-sampled and used in this study. This seaweed had bromoform concentration of 6.55 mg/g, which was marginally lower than our study and may be due to variation in the collection, sampling or analysis techniques. Despite the lower bromoform concentration in the seaweed and using 0.20% inclusion rate of *A. taxiformis* on OM basis, the CH_4_ yield was reduced by up to 98% in Brangus feedlot steers. The diet used by Kinley et al. (2020) included 30.6% NDF, which was similar to our high fiber diet.

The greater efficacy of *A. taxiformis* in that study could be due to collective feed formulation differences such as the energy dense component of barley versus corn, which is typical of Australian and American feedlots, respectively. Additionally, it could be due to beneficial interaction with the ionophore, monensin, that was used in the Australian study. Monensin has not been used in any other feed formulation in other *in vivo* studies with the inclusion of *Asparagopsis* species. Use of monensin in diets has shown to decrease CH_4_ yields by up to 6% in feedlot steers while also having an enhanced effect in diets containing greater NDF levels (Appuhamy et al., 2013). A potential enhancing interaction of the seaweed with monensin is of great interest and further investigation will elucidate this potential that could have significant beneficial economic and environmental impact for formulated feeding systems that use monensin.

### 4.2 Enteric hydrogen and carbon dioxide emissions

Increases in H_2_ yield have typically been recorded when anti-methanogenic feed additives are used, and with the addition of *Asparagopsis* species in dairy cattle (1.25 to 3.75 fold; Roque et al., 2019b) and Brangus feedlot steers (3.8-17.0 fold; Kinley et al., 2020). Similar increases in H_2_ yield have been reported in feed additives that reduce enteric CH_4_ emissions targeting methanogens. For example, in lactating dairy cows supplemented with 3-NOP, H_2_ yield increased 23 – 71 fold (Hristov et al., 2015). Bromochloromethane (BCM) fed to goats increased H_2_ (mmol/head per day) 5 – 35 fold, while chloroform fed to Brahman steers increased H_2_ yield 316 fold (Mitsumori et al., 2012; Martinez-Fernandez et al., 2016). Although feeding *Asparagopsis spp*. increased overall H_2_ yield (Figure 4), the magnitude was considerably lower (1.25 to 17 fold) compared to alternative CH_4_ reducing feed additives (5 to 316 fold), with similar levels of reductions in CH_4_. This indicates that there may be a redirection of H_2_ molecules that would otherwise be utilized through the formation of CH_4_ and redirected into different pathways that could be beneficial to the animal. For example, increased propionate to acetate concentrations have been recorded in *in vitro* studies using *A. taxiformis* (Machado et al., 2016a; Roque et al., 2019a) during inhibition of CH_4_ which could indicate that some of the excess H_2_ is being utilized for propionate production. Consistent with this theory, similar results have been reported using other CH_4_ analogues such as BCM (Denmen et al., 2015) and chloroform, (Martinez-Fernandez et al., 2016) both of which showed increases in propionate production. In general, significant reduction of CH_4_ production in the rumen without detriment to rumen function is typically associated with reduction of acetate, increase in propionate, and reduction of the acetate:propionate ratio (Kinley et al., 2020).

**FIGURE 4.**
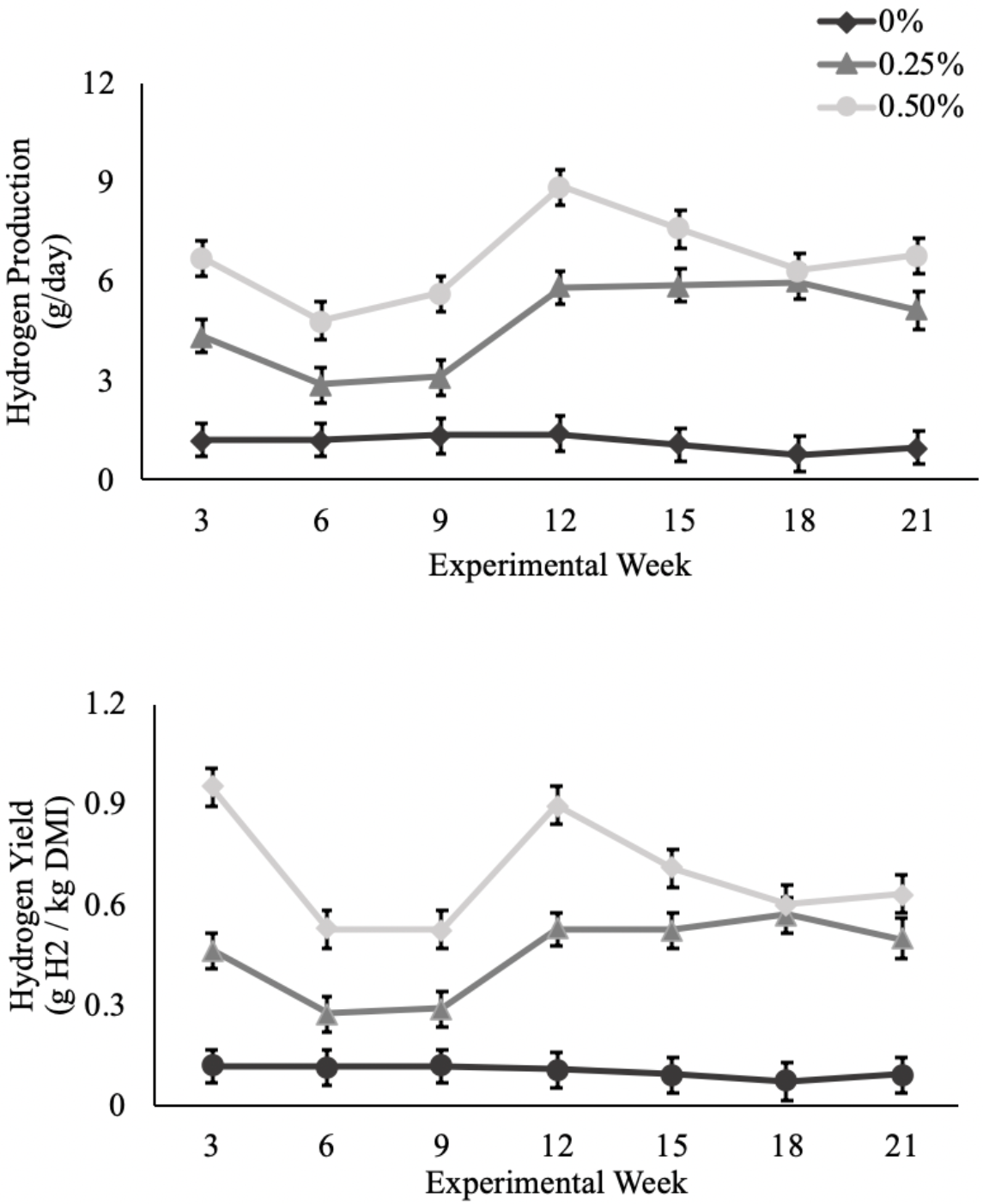
Hydrogen production [g H_2_/day] (A) and Hydrogen yield [g H_2_/kg DMI] (B) from beef steers supplemented with *Asparagopsis taxiformis* at 0%, 0.25%, and 0.5% of basal total mixed ration on an organic matter basis during the 21 week experimental period. Data points are treatment means for each gas collection timepoint and error bars represent standard errors.

In contrast to the lactating dairy cattle study in which CO_2_ yield increased significantly in dairy cattle fed 1% *A. armata* (Roque et al., 2019b), there was no significant difference in CO_2_ emissions or yield in the current study. This could be due to the relationship between the amount of seaweed fed and DMI intake.

### 4.3 Animal production parameters

Dry matter intake reductions observed in this study were consistent with previous studies in lactating dairy cows where decreases in DMI were found to be 10.7 and 37.9% at 0.50 and 1.0% inclusion rate of *A. armata* (Roque et al., 2019b), respectively. Decreases in DMI have also been reported in cattle fed other anti-methanogenic feed additives in a linear dose-response fashion. For example, Tomkins et al. (2009) reported 3 to 19% reductions in DMI in steers supplemented with BCM at dosages between 0.15 and 0.60 g/100 kg live weight. Additionally, Martinez-Fernandez (2016) found 1.7 to 15% reductions in DMI when feeding steers chloroform at dosages between 1 to 2.6 g/100g liveweight. In contrast, Kinley et al. (2020) reported no significant differences at the highest *A. taxiformis* level of 0.20%. However, the inclusion level was less than our study’s lowest inclusion rate, so based on previous experiment’s observation of reduced DMI in a dose-response manner (Roque et al., 2019b), it was expected to have lower effect on DMI. Decreases in DMI are normally associated with lower productivity due to lower levels of nutrients and dietary energy consumed. However, there was no significant difference in ADG between steers in the HD treatment and Control (average 1.56 kg/day) groups despite consuming 14% less feed. The results were in agreement with a previous study (Roque et al., 2019b), in which milk production was not compromised at a 0.5% OM inclusion level despite reductions in DMI. The FCE (ADG/DMI) increased significantly in HD treatment group, suggesting that inclusion rates of *A. taxiformis* at 0.5% improves overall feed efficiency in growing beef steers. Since a large proportion of on farm costs is the purchase of feed, an improved feed efficiency is particularly exciting for producers to decrease feed costs while also producing the same amount of total weight gains. Total gains were between 224 kg (LD) to 236 kg (HD) combined with an average cost differential of ~$0.18 USD/kg gain (LD) and ~$0.37 USD/kg gain (HD). A producer finishing 1000 head of beef cattle has the potential to reduce feed costs by $40,320 (LD) to $87,320 (HD) depending on seaweed dosage. While the CPG in this study were not statistically significant, this may be due to low animal numbers in each treatment and warrants further investigation on a larger feedlot setting to reduce animal variability.

### 4.4 Bromoform and iodine residues

Bromoform is considered to be the active ingredient responsible for CH_4_ reduction when fed to cattle (Machado et al., 2016b). However, high levels of bromoform is considered to be hazardous for humans and mice. While bromoform intake limits are yet to be defined for cattle specifically, the United States Environmental Protection Agency (2017) has suggested a reference dose for bromoform, an estimated level of daily oral exposure without negative effects, to be 0.02 mg/kg BW/day. It is essential that food products from livestock consuming the seaweed are confirmed as safe for consumption and that bromoform residues are not transferred to the edible tissues and offal of bovines at levels detrimental to food safety. Previous studies have demonstrated that bromoform was not detectable in the kidney, muscle, fat deposits, blood, feces, and milk in either sheep (Li et al., 2018), dairy cows (Roque et al., 2019b), or feedlot steers (Kinley et al., 2020). Strip loin and liver samples from steers were collected and in agreement with previous studies, no bromoform was detected in this study.

The National Academies of Sciences, Engineering, and Medicine (2016) recommendations for daily iodine intake in growing beef cattle is 0.5 ug/g DMI and maximum tolerable limit is 50 ug/g DMI. In this study, recommended daily iodine intake levels were 5.2 mg/day and 4.85 mg/day and maximum limits are 521 mg/day and 485 mg/day for LD and HD treatment groups, respectively. The iodine level in the *A. taxiformis* fed in the current study contained 2.27 mg/g, therefore, maximum daily intake of seaweed iodine was 106 – 127 mg/day and 173 – 225 mg/day for the LD and HD treatment groups, respectively. While these levels do not exceed maximum tolerable limits, they exceed daily iodine intake recommendations, therefore it was appropriate to test for iodine residue levels in meat used for human consumption. The US Food and Nutrition Board of the National Academy of Sciences has set a tolerable upper intake level (UL) for human consumption of foods, which is defined as the highest level of daily intake that poses no adverse health effects (Trumbo et al., 2001). The iodine UL ranges between 200 ug/day to 1,100 ug/day depending on age, gender, and lactation demographics. Strip loins tested for iodine residues had levels of 0.08 and 0.15 ug/g from steers in treatments LD and HD, respectively. These iodine residues are far under the UL limits for human consumption. For example, UL for a person under 3 years of age is 200 ug/day meaning that this person would have to consume more than 2,500 g/day and 1,330 g/day of meat from a LD and HD steers, respectively, to reach the UL. An adult over the age of 18 has an UL of 1,100 ug/day and would have to consume more than 13.8 kg/day and 7.3 kg/day of meat from a LD and HD steers, respectively, to reach their UL of iodine intake. At the inclusion levels and iodine concentration of *A. taxiformis* used in this study the margin of safety is extremely high and the likelihood of iodine toxicity from consuming the meat is extremely low.

### 4.5 Carcass quality parameters

Marbling scores ranged from 410 – 810 while all carcasses, regardless of treatment, graded as either choice or prime. The value placed on tenderness in the marketplace is high and has even been found that consumers are likely to pay premiums for more tender beef (Miller et. al, 2001). Rodriquez-Herrera et al (2016) found that cattle supplemented with a high amount of *Schizochytrium spp.* (30 g/kg of feed intake) had improved tenderness compared to the control treatment and a low dose treatment group. Conversely, Phelps et al (2016) reported that supplementation of *Schizochytrium spp.* in cattle at 0 to 150 g/day per animal, a lower inclusion rate than Rodriquez-Herrera et al (2016), did not have a significant impact on objective tenderness. While no significant differences in meat tenderness were observed for either WBSF or SSF values among the three treatment groups in this study, it may be due to low inclusions rates and warrants further investigation. If the macroalgae *A. taxiformis* has the potential to perform similarly to the microalgae *Schizochytrium spp.*, this could increase consumers’ preferences for algae-fed beef while also providing producers with a tenderness premium for their product.

The current study showed no statistical differences in consumer sensory evaluation or preference toward steaks from any of the treatment groups in agreement with Kinley et al. (2020) in which steaks from beef cattle fed *A. taxiformis* were evaluated for taste, tenderness, juiciness, or overall flavor. Another study demonstrated that trained panelists were unable to detect a difference in tenderness in beef steaks from cattle fed up to 150 g of *Schizochytrium spp.* per day (Phelps et al, 2016). Conversely, some studies reported that the supplementation of *Schyzochytrium* fed at a 3.89% inclusion rate of DMI in lamb diets and up to 30 g/kg of feed intake to cattle resulted in meat samples as having a “seaweed” or “fishy” flavor in which the authors attributed to increased docosahexaenoic acid levels (Rodriguez-Herrera et. al, 2018; Urrutia et. al., 2016). However, these two studies feed a substantially greater amount of seaweed in their diets than the current study, which could be the reason for the alteration in overall taste. The current study indicates that the supplementation of *A. taxiformis* at 0.25% or 0.5% to cattle does not significantly impact overall meat quality nor alter the sensory properties of the steaks.

In summary, this study showed that the use of *A. taxiformis* supplemented to beef cattle diets reduced enteric CH_4_ emissions for a duration of 21 weeks without any loss in efficacy. The efficacy was highly correlated with the proportion of NDF in the diet. Additionally, *A. taxiformis* has no measurable residual effect in the product and did not alter meat quality or sensory properties. Importantly, the use of *A. taxiformis* impacts DMI and not ADG, therefore increasing overall feed efficiency (FCE) in growing beef steers. There may also be potential to reduce the cost of production per kg of weight gain. These feed cost reductions in combination with significantly reduced CH_4_ emissions have a potential to transform beef production into a more financially and environmentally sustainable and product efficient industry.

## ACKNOWLEDGEMENTS

This research received financial support from Elm Innovations, the David and Lucile Packard Foundation and the Grantham Foundation. The authors acknowledge Meat and Livestock Australia, James Cook University and CSIRO for the supply of *A. taxiformis* used in the trial. We are grateful to undergraduate interns: A. Neveu, A. Wilson, A. Yiao, B. Wong, C. Chow, C. Martinez, C. Mielke, D. Maqueda, E. Anderson, J. DeGuzman, J. Fang, J. Infante, J. Jordan, K. Allchin, K. Garcia, K. Martin, L. Arkangel, M. Cervantes, M. Venegas, M. Zack, P. Nguyen, P. Petschl, S. Calderon, S. Leal, S. Lee, T. Lee, and V. Escobar that participated in the trial. We appreciate Dr. Craig Burnell and Steve Archer (Bigelow Labs in East Boothbay, ME, USA) for developing methods to measure bromoform concentration in *Asparagopsis taxiformis*, meat, liver, and feces collected in this study.

## REFERENCES

Abecia, L., Toral, P.G., Martín-García, A.I., Martínez, G., Tomkins, N.W., Molina-Alcaide, E., Newbold, C.J. and Yáñez-Ruiz, D.R. (2012). Effect of bromochloromethane on methane emission, rumen fermentation pattern, milk yield, and fatty acid profile in lactating dairy goats. Journal of Dairy Science, 95(4), 2027–2036. https://doi.org/10.3168/jds.2011-4831

AMSA. (2016). Research guidelines for cookery, sensory evaluation, and instrumental tenderness measurements of meat. American Meat Science Association (AMSA): 1–105. www.meatscience.org

Appuhamy, J.R.N., Strathe, A.B., Jayasundara, S., Wagner-Riddle, C., Dijkstra, J., France, J. and Kebreab, E. (2013). Anti-methanogenic effects of monensin in dairy and beef cattle: A meta-analysis. Journal of Dairy Science, 96(8), 5161–5173. https://doi.org/10.3168/jds.2012-5923

Bannink, A., France, J., Lopez, S., Gerrits, W.J.J., Kebreab, E., Tamminga, S. and Dijkstra, J. (2008). Modelling the implications of feeding strategy on rumen fermentation and functioning of the rumen wall. Animal Feed Science and Technology, 143(1-4), 3–26. https://doi.org/10.1016/j.anifeedsci.2007.05.002

Bannink, A., Kogut, J., Dijkstra, J., France, J., Kebreab, E., Van Vuuren, A.M. and Tamminga, S. (2006). Estimation of the stoichiometry of volatile fatty acid production in the rumen of lactating cows. Journal of Theoretical Biology, 238(1), 6–51. https://doi.org/10.1016/j.jtbi.2005.05.026

Blaxter, K.L. and Clapperton, J.L. (1965). Prediction of the amount of methane produced by ruminants. British Journal of nutrition, 19(1), 511–522. https://doi.org/10.1079/BJN19650046

Denman, S.E., Fernandez, G.M., Shinkai, T., Mitsumori, M., and McSweeney, C.S. (2015). Metagenomic analysis of the rumen microbial community following inhibition of methane formation by a halogenated methane analog. Frontiers in Microbiology, 6, 1087. https://doi.org/10.3389/fmicb.2015.01087

Dijkstra, J., Bannink, A., France, J., Kebreab, E., and van Gastelen, S. (2018). Short communication: antimethanogenic effects of 3-nitrooxypropanol depend on supplementation dose, dietary fiber content, and cattle type. Journal of Dairy Science. 101, 9041e9047. https://doi.org/10.3168/jds.2018-14456.

Duin, E.C., Wagner, T., Shima, S., Prakash, D., Cronin, B., Yáñez-Ruiz, D.R., Duval, S., Rümbeli, R., Stemmler, R.T., Thauer, R.K., and Kindermann, M. (2016). Mode of action uncovered for the specific reduction of methane emissions from ruminants by the small molecule 3-nitrooxypropanol. Proceedings of the National Academy of Sciences United States of America. 113, 6172e6177. https://doi.org/10.1073/pnas.1600298113.

Ermler, U., Grabarse, W., Shima, S., Goubeaud, M., and Thauer, R.K. (1997). Crystal structure of methyl-coenzyme M reductase: the key enzyme of biological methane formation. Science. 278 (5342), 1457e1462. https://10.1126/science.278.5342.1457.

Gerber, P.J., Steinfeld, H., Henderson, B., Mottet, A., Opio, C., Dijkman, J., Falcucci, A., and Tempio, G. (2013). Tackling Climate Change through Livestock: a Global Assessment of Emissions and Mitigation Opportunities. FAO, Rome. http://www.fao.org/docrep/018/i3437e/i3437e00.htm.

Hungate, R. (1966). Chapter VI - Quantities of Carbohydrate Fermentation Products: The Rumen and its Microbes. Academic Press, 245–280, https://doi.org/10.1016/B978-1-4832-3308-6.50009-7.

Hristov, A.N., Oh, J., Firkins, J.L., Dijkstra, J., Kebreab, E., Waghorn, G., Makkar, H. P. S., Adesogan, A. T., Yang, W., Lee, C., Gerber, P. J., Henderson, B. and Tricarico, J. M. (2013). Mitigation of methane and nitrous oxide emissions from animal operations: I. A review of enteric methane mitigation options. Journal of Animal Science, 91, 5045–5069.

Hristov, A.N., Oh, J., Giallongo, F., Frederick, T.W., Harper, M.T., Weeks, H.L., Branco, A.F., Moate, P.J., Deighton, M.H., Williams, S.R.O. and Kindermann, M. (2015). An inhibitor persistently decreased enteric methane emission from dairy cows with no negative effect on milk production. Proceedings of the National Academy of Sciences, 112(34), 10663–10668. https://doi.org/10.1073/pnas.1504124112

Janssen, PH. 2010. Influence of hydrogen on rumen methane formation and fermentation balances through microbial growth kinetics and fermentation thermodynamics. Animal Feed Science and Technology, (160), 1–22.

Johnson, E. D., Wood, A. S., Stone, J. B., and Moran Jr, E. T. (1972). Some effects of methane inhibition in ruminants (steers). Canadian Journal of Animal Science, 52(4), 703–712. https://doi.org/10.4141/cjas72-083

Johnson, K. A., & Johnson, D. E. (1995). Methane emissions from cattle. Journal of Animal Science, 73(8), 2483–2492.

Killinger, K. M., Calkins, C. R., Umberger, W. J., Feuz, D. M., and Eskridge, K. M. (2004). Consumer sensory acceptance and value for beef steaks of similar tenderness, but differing in marbling level. Journal of Animal Science, 82(11), 3294–3301. https://doi.org/10.2527/2004.82113294x

Kinley, R.D., de Nys, R., Vucko, M.J., Machado, L., Tomkins, N.W. (2016a). The red macroalgae Asparagopsis taxiformis is a potent natural antimethanogenic that reduces methane production during in vitro fermentation with rumen fluid. Animal Production Science, (56), 282e289. https://doi.org/10.1071/AN15576.

Kinley, R.D., Vucko, M.J., Machado, L., Tomkins, N.W. (2016b). In vitro evaluation of the antimethanogenic potency and effects on fermentation of individual and combinations of marine macroalgae. American Journal of Plant Sciences, (7), 2038e2054. https://doi.org/10.4236/ajps.2016.714184.

Kinley, R.D., Martinez-Fernandez, G., Matthews, M.K., de Nys, R., Magnusson, M. and Tomkins, N.W. (2020). Mitigating the carbon footprint and improving productivity of ruminant livestock agriculture using a red seaweed. Journal of Cleaner Production, 59, 120836. https://doi.org/10.1016/j.jclepro.2020.120836

Knight, T., Ronimus, R.S., Dey, D., Tootill, C., Naylor, G., Evans, P., Molano, G., Smith, A., Tavendale, M., Pinares-Patino, C.S. and Clark, H. (2011). Chloroform decreases rumen methanogenesis and methanogen populations without altering rumen function in cattle. Animal Feed Science and Technology, 166, 101–112. https://doi.org/10.1016/j.anifeedsci.2011.04.059

Li, X., Norman, H.C., Kinley, R.D., Laurence, M., Wilmot, M., Bender, H., de Nys, R. and Tomkins, N. (2018). *Asparagopsis taxiformis* decreases enteric methane production from sheep. Animal Production Science, 58(4), 681–688. https://doi.org/10.1071/AN15883

Liu, H., Wang, J., Wang, A., and Chen, J. (2011). Chemical inhibitors of methanogenesis and putative applications. Applied Microbiology and Biotechnology, 89, 1333e1340. https://doi.org/10.1007/s00253-010-3066-5.

Machado, L., Magnusson, M., Paul, N.A., de Nys, R., and Tomkins, N. (2014). Effects of Marine and Freshwater Macroalgae on In-Vitro Total Gas and Methane Production. PloS One, (9)1. https://doi.org/10.1371/journal.pone.0085289

Machado, L., Magnusson, M., Paul, N.A., Kinley, R., de Nys, R., and Tomkins, N. (2016a). Dose-response effects of Asparagopsis taxiformis and Oedogonium sp. on in-vitro fermentation and methane production. Journal of Applied Phycology. 28,1443–1452. https://doi.org/10.1007/s10811-015-0639-9

Machado, L., Magnusson, M., Paul, N.A., Kinley, R., de Nys, R., and Tomkins, N. (2016b). Identification of bioactives from the red seaweed Asparagopsis taxiformis that promote antimethanogenic activity in-vitro. Journal of Applied Phycology. 28, 3117–3126. https://doi.org/10.1007/s10811-016-0830-7

Machado, L., Tomkins, N., Magnusson, M., Midgley, D., deNyes, R., and Rosewarne, C. (2018). In vitro response of rumen microbiota to the antimethanogenic red macroalga Asparagopsis taxiformis. Microbial Ecology. 75, 811–818. https://doi.org/10.1007/s00248-017-1086-8

Martinez-Fernandez, G., Denman, S. E., Yang, C. L., Cheung, J. E., Mitsumori, M., and Mcsweeney, C. S. (2016). Methane inhibition alters the microbial community, hydrogen flow, and fermentation response in the rumen of cattle. Frontiers in Microbiology, 7(1122), https://doi.org/10.3389/fmicb.2016.01122

Mitsumori, M., Shinkai, T., Takenaka, A., Enishi, O., Higuchi, K., Kobayashi, Y., Nonaka, I., Asanuma, N., Denman, S.E. and McSweeney, C.S. (2012). Responses in digestion, rumen fermentation and microbial populations to inhibition of methane formation by a halogenated methane analogue. British Journal of Nutrition, 108(3), 482–491. https://doi.org/10.1017/S0007114511005794

Miller, M. F., Carr, M. A., Ramsey, C. B., Crockett, K. L., & Hoover, L. C. (2001). Consumer thresholds for establishing the value of beef tenderness. Journal of animal science, 79(12), 3062–3068. https://doi.org/10.2527/2001.79123062x

Moraes, L.E., Strathe, A.B., Fadel, J.G., Casper, D.P., and Kebreab, E. (2014). Prediction of enteric methane emissions from cattle. Global Change Biology. 20 (7), 2140e2148. https://doi.org/10.1111/gcb.12471.

Niu, M., Kebreab, E., Hristov, AN., et al. (2018). Prediction of enteric methane production, yield, and intensity in dairy cattle using an intercontinental database. Global Change Biology. 24: 3368–3389. https://doi.org/10.1111/gcb.14094

Paul, N.A., Cole, L., de Nys, R., and Steinberg, P.D. (2006). Ultrastructure of the gland cells of the red alga Asparagopsis armata (Bonnemaisoniaceae). Journal of Phycology. 42, 637–645. https://doi.org/10.1111/j.1529-8817.2006.00226.x.

Phelps, K. J., Drouillard, J. S., O’Quinn, T. G., Burnett, D. D., Blackmon, T. L., Axman, J. E., … and Gonzalez, J. M. (2016). Feeding microalgae meal (All-G Rich; CCAP 4087/2) to beef heifers. I: Effects on longissimus lumborum steak color and palatibility. Journal of Animal Science, 94(9), 4016–4029. https://doi.org/10.2527/jas.2016-0487

Rodriguez-Herrera, M., Khatri, Y., Marsh, S. P., Posri, W., & Sinclair, L. A. (2018). Feeding microalgae at a high level to finishing heifers increases the long-chain n-3 fatty acid composition of beef with only small effects on the sensory quality. International Journal of Food Science & Technology, 53(6), 1405–1413. https://doi.org/10.1111/ijfs.13718

Roque, B.M., Brooke, C.G., Ladau, J., Polley, T., Marsh, L.J., Najafi, N., Pandey, P., Singh, L., Kinley, R., Salwen, J.K., Eloe-Fadrosh, E., Kebreab, E., and Hess, M. (2019a). Effect of the macroalgae *Asparagopsis taxiformis* on methane production and rumen microbiome assemblage. Animal Microbiome, 1(1), 3. https://doi.org/10.1186/s42523-019-0005-3

Roque, B.M., Salwen, J.K., Kinley, R. and Kebreab, E. (2019b). Inclusion of *Asparagopsis armata* in lactating dairy cows’ diet reduces enteric methane emission by over 50 percent. Journal of Cleaner Production, 234, 132–138. https://doi.org/10.1016/j.jclepro.2019.06.193

Rumpler, W.V., Johnson, D.E. and Bates, D.B. (1986). The effect of high dietary cation concentration on methanogenesis by steers fed diets with and without ionophores. Journal of Animal Science, 62(6), 1737–1741. https://doi.org/10.2527/jas1986.6261737x

Russell, J. B., & Wallace, R. J. (1997). Energy-yielding and energy-consuming reactions: The rumen microbial ecosystem Springer Dordrecht, Chicago. 246–282.

Smith, E.L., Mervyn, L., Johnson, A.W. and Shaw, N. (1962). Partial synthesis of vitamin B12 coenzyme and analogues. Nature, 194(4834), 1175–1175. https://doi.org/10.1038/1941175a0

Tomkins, N.W., Colegate, S.M. and Hunter, R.A. (2009). A bromochloromethane formulation reduces enteric methanogenesis in cattle fed grain-based diets. Animal Production Science, 49(12), 1053–1058. https://doi.org/10.1071/EA08223

Trumbo, P., Yates, A.A., Schlicker, S. and Poos, M. (2001). Dietary reference intakes: vitamin A, vitamin K, arsenic, boron, chromium, copper, iodine, iron, manganese, molybdenum, nickel, silicon, vanadium, and zinc. Journal of the Academy of Nutrition and Dietetics, 101(3), 294. https://doi.org/10.1016/S0002-8223(01)00078-5

United States Environmental Protection Agency. (2017). Integrated Risk Information System (IRIS) on Bromoform. National Center for Environmental Assessment, Office of Research and Development, Washington, D.C. https://cfpub.epa.gov/ncea/iris

Urrutia, O., Mendizabal, J. A., Insausti, K., Soret, B., Purroy, A., & Arana, A. (2016). Effects of addition of linseed and marine algae to the diet on adipose tissue development, fatty acid profile, lipogenic gene expression, and meat quality in lambs. PloS One, 11(6). https://doi.org/10.1371/journal.pone.0156765

Van Soest, P. J. (1994). Nutritional ecology of the ruminant. Cornell University Press.

Wood, J. M., Kennedy, F. S., and Wolfe, R. S. (1968). Reaction of multihalogenated hydrocarbons with free and bound reduced vitamin B12. Biochemistry, 7(5), 1707–1713. https://doi.org/10.1021/bi00845a013

